# Integer programming framework for pangenome-based genome inference

**DOI:** 10.1101/2024.10.27.620212

**Authors:** Ghanshyam Chandra, Md Helal Hossen, Stephan Scholz, Alexander T Dilthey, Daniel Gibney, Chirag Jain

**Affiliations:** Department of Computational and Data Sciences, Indian Institute of Science, Bangalore KA 560012, India; Department of Computer Science, The University of Texas at Dallas, TX 75080, USA; Institute of Medical Microbiology and Hospital Hygiene, Heinrich Heine University Düsseldorf, Düsseldorf, Germany; Center for Digital Medicine, Heinrich Heine University Düsseldorf, Düsseldorf, Germany

## Abstract

Affordable genotyping methods are essential in genomics. Commonly used genotyping methods primarily support single nucleotide variants and short indels but neglect structural variants. Additionally, accuracy of read alignments to a reference genome is unreliable in highly polymorphic and repetitive regions, further impacting genotyping performance. Recent works highlight the advantage of haplotype-resolved pangenome graphs in addressing these challenges. Building on these developments, we propose a rigorous alignment-free genotyping framework. Our formulation seeks a path through the pangenome graph that maximizes the matches between the path and substrings of sequencing reads (e.g., *k*-mers) while minimizing recombination events (haplotype switches) along the path. We prove that this problem is NP-Hard and develop efficient integer-programming solutions. We benchmarked the algorithm using downsampled short-read datasets from homozygous human cell lines with coverage ranging from 0.1× to 10×. Our algorithm accurately estimates complete major histocompatibility complex (MHC) haplotype sequences with small edit distances from the ground-truth sequences, providing a significant advantage over existing methods on low-coverage inputs. Although our algorithm is designed for haploid samples, we discuss future extensions to diploid samples.

**Implementation:** https://github.com/at-cg/PHI

## 1 Introduction

Many initiatives are in progress for building haplotype-resolved pangenome references of human and non-human species [22,11,36]. Among many applications, pangenome graphs can enable cost-effective genotyping and imputation of a wide spectrum of variant classes beyond single nucleotide polymorphisms (SNPs) and short indels [13]. Pangenome graphs represent sequence alignment of high-quality fully-phased genome assemblies of individuals from diverse populations [1]. A pangenome graph can be represented as either cyclic or acyclic directed graph where the vertices are labeled with sequences. Paths in this graph spell the reference haplotype sequences and their recombinations. The graph-based representation is flexible enough to incorporate single-nucleotide polymorphisms (SNPs), indels (short insertions and deletions), large structural variants (SVs), nested variants, gene absence/presence, etc. [4].

Recent works propose the use of pangenome references to improve genotyping accuracy from short-read sequencing data [9,14,12,18,2,10,33,25]. Especially for SVs, these methods are an effective alternative to the conventional genotyping methods that are based on aligning reads to a single reference because short-read alignments can be inaccurate for the reads originating from SVs [23,8]. Methods such as PRG [6], Pangenie [9], and KAGE [12], utilize *k*-mer statistics to infer paths in the graph that correspond to the target genome. These methods compare the *k*-mers surrounding a variant site in the graph with the *k*-mer counts in the sequencing data to calculate likelihoods of reference and alternative alleles. Pangenie and KAGE also use the long-range haplotype information available in the haplotype-resolved pangenome references. The other approach used in methods such as Giraffe [34] and Graphtyper [10] involves aligning reads to a pangenome graph.

There have been efforts on improving the accuracy of read alignments to pangenome graphs as well. A large combinatorial search space in terms of the number of candidate paths in a pangenome graph increases ambiguity during read alignment. This issue has motivated methods that either impute a personalized reference genome [38], sample variants [29,17,37] to obtain a smaller graph, or prioritize the use of reference haplotypes in the graph during alignment [3,34,26]. Our previous work proposed haplotype-aware sequence alignment to graphs by introducing penalties for haplotype switches in an alignment [3]. A recent feature added to VG allows sampling of reference haplotypes and their recombinations from the graph that are most relevant to the target genome using a *k*-mer-based greedy heuristic [35].

Low-coverage sequencing, combined with genotyping and phasing, is a cost-effective approach to conduct large-scale genetic studies [31,5,20,24]. In this paper, we develop a rigorous formulation and algorithms for genotyping using pangenome references. Our framework is also applicable to low-coverage short-read sequencing data (coverage 0.1−1×). Following the standard Li and Stephens model [21], we view the target genome as an imperfect mosaic of the reference haplotypes. Our contributions are as following.

– We introduce a novel problem formulation to estimate the complete haplotype sequence of a haploid genome by determining an appropriate path in the pangenome graph. The objective is to maximize the number of shared substrings (e.g., *k*-mers or minimizers) between the sequencing data and the sequence spelled by the path. We permit recombinations in the path, subject to a fixed penalty per recombination. We refer to this problem as *Path Inference Problem* (formally defined in Section 2).
– We prove that the Path Inference Problem is NP-hard, even when restricted to binary alphabets.
– To solve this problem, we develop two integer-programming solutions which involve linear and quadratic constraints, respectively. The two solutions involve a tradeoff between runtime and memory usage.
– We demonstrate the utility of this framework by testing it on downsampled short-read datasets from five human haploid cell lines (coverage 0.1 − 10×). For these five samples, complete major histocompatibility complex (MHC) haplotype sequences have been previously determined using long-read assembly [16]. As our pangenome reference, we used a haplotype-resolved pangenome directed acyclic graph (DAG) of 49 MHC haplotype sequences [19]. We chose MHC region for evaluation because this is the most polymorphic and gene-rich region of the human genome [7]. The length of this region is about 5 Mbp.
– Using datasets with 0.1× coverage, our algorithm outputs MHC sequences that are up to 99.96% identical to the ground-truth sequences. It compares favorably to the existing methods.

## 2 Notations and Problem Formulation

Let *G*(*V, E, σ*, ℋ) denote a directed acyclic graph (DAG) representing a haplotype-resolved pangenome reference. Function *σ* assigns a string label over alphabet *Σ* = {*A, C, G, T*} to each vertex. A path (*u*_1_, *u*_2_, …, *u*_*n*_) in *G* spells string *σ*(*u*_1_) · *σ*(*u*_2_) · · · *σ*(*u*_*n*_), where *s*_1_ · *s*_2_ denotes the concatenation of strings *s*_1_ and *s*_2_. ℋ = {*h*_1_, *h*_2_, …, *h*_|ℋ|_}denotes a set of paths in *G* such that each of these paths spells a reference haplotype sequence used in the pangenome reference. We refer to these paths as haplotype paths. We assume that each haplotype path is described by an array, i.e., *h*_*i*_[1] is the first vertex in *h*_*i*_, *h*_*i*_[2] is the second vertex in *h*_*i*_, etc. The length of a haplotype path *h*_*i*_, that is, the count of vertices in *h*_*i*_ is denoted as |*h*_*i*_|. The set of haplotype paths covering vertex *ν* ∈ *V* is denoted as *haps*(*ν*). We assume that, for each edge (*u, ν*) ∈ *E*, there exists a haplotype path *h*_*i*_ ∈ ℋ such that *u* and *ν* are consecutive vertices in *h*_*i*_. In other words, each edge is supported by at least one haplotype path.

### Definition 1 (Inferred Path).

*An inferred path of* 𝒫 *length n is represented as an ordered set* (*a*_1_, *a*_2_, …, *a*_*n*_), *where each a*_*i*_ *is a two tuple* (*u, h*) *such that u* ∈ *V, h* ∈ *haps*(*u*), *and* (*a*_*i*_.*u, a*_*i*+1_.*u*) ∈ *E for all i* ∈ [1, *n*). *Furthermore, if a*_*i*_.*h* = *a*_*i*+1_.*h, then a*_*i*_.*u and a*_*i*+1_.*u should be consecutive vertices in haplotype path a*_*i*_.*h*.

In an inferred path, we keep track of the haplotype path indices alongside vertex indices (Figure 1). We say a *recombination*, or a haplotype switch, occurs between two consecutive vertices *a*_*i*_.*u* and *a*_*i*+1_.*u* in 𝒫 if *a*_*i*_.*h* ≠ *a*_*i*+1_.*h*. We use *γ*(𝒫) to denote the count of recombinations in 𝒫. With a mild abuse of notation, we denote the string spelled by 𝒫 as *σ*(𝒫).

**Fig. 1:**
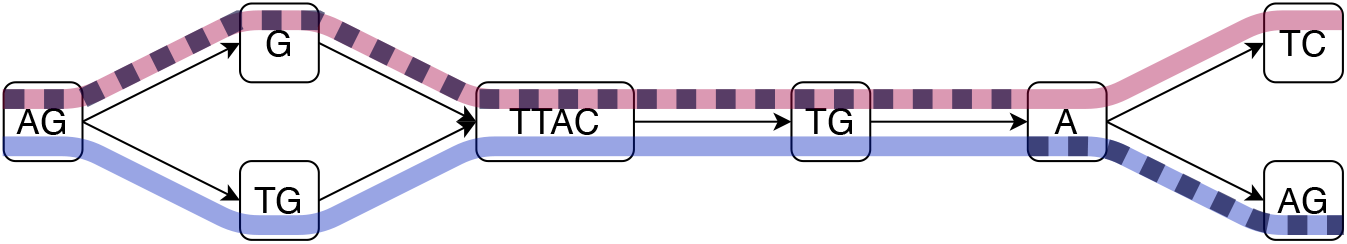
A simple illustration of an haplotype-resolved pangenome graph with two haplotype paths highlighted in pink and blue colors. An inferred path with a single recombination is shown as a dashed line.

### Problem 1

(Path Inference Problem).

– **Input:** A haplotype-resolved pangenome DAG *G* = (*V, E, σ*, ℋ), a set of strings 𝒮 from the target genome, and a non-negative integer *c* indicating recombination penalty.
– **Output:** An inferred path 𝒫such that

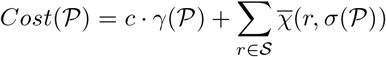

is minimized, where 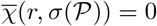 if string *r* occurs as a substring of string *σ*(𝒫) and 1 otherwise.

The intuition behind our formulation is to maximize the number of string matches along the inferred path while minimizing the number of recombinations. This approach yields an inferred path that incorporates the majority of strings from 𝒮 as a substring with a finite number of recombinations, constrained by a recombination penalty *c*. Set 𝒮 can be set of either *k*-mers or minimizers observed in the sequencing reads.

## 3 Computational Complexity

### Theorem 1.

*Problem 1 is NP-hard. This holds for any value of c* = |*V* |^*Θ*(1)^ *and even when Σ* = {0, 1}.

We begin with an instance *G*_*H*_ (*V*_*H*_, *E*_*H*_) of the Hamiltonian Path Problem. Let *V*_*H*_ = {*u*_1_, …, *u*_*n*_}. We first create a graph *G*′ = (*V* ′, *E*′) where

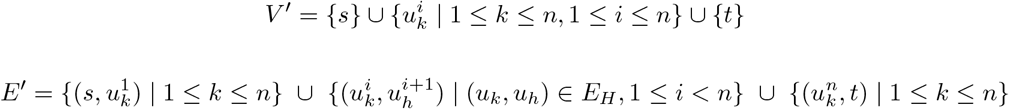

For 1 ≤ *x* ≤ *n* + 2(*c*(*n* + 1) + 1, let bin(*x*) be standard binary encoding of *x* using *b* = ⌈log_2_(*n* + 2(*c*(*n* + 1) + 1))⌉ + 1 bits. We assign the vertex labels

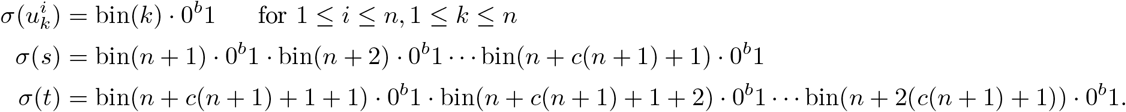

We create a distinct haplotype path for each edge that supports only that edge. We define the set of strings 𝒮 = {bin(1)·0^*b*^1, bin(2)·0^*b*^1, …, bin(*n* + 2(*c*(*n* + 1) + 1))· 0^*b*^1}. See Figure 1 in Appendix for a small worked example. The reduction presented above clearly runs in polynomial time for *c* = |*V*| ^*Θ*(1)^. Combined with Lemmas 1 and 2, Theorem 1 follows.

### Lemma 1.

*If G*_*H*_ *contains a Hamiltonian path, then G*′ *has an inferred path* 𝒫*with Cost*(𝒫) = *c* · (*n* + 1)

*Proof*. Let *u*_*i*_, …, *u*_*i*_ be a Hamiltonian path in *G*_*H*_. We take as our inferred path 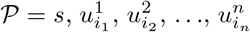, *t*. As every edge has its own corresponding haplotype, the number of recombinations is *n* + 1. Furthermore, since 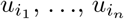 is a Hamiltonian path and *s* and *t* are included in the inferred path, all strings in 𝒮 occur in *σ*(𝒫). Hence, the total cost is *c* · (*n* + 1). □

### Lemma 2.

*If G*′ *has an inferred path* 𝒫*with Cost*(𝒫) ≤ *c* · (*n* + 1), *then G*_*H*_ *has a Hamiltonian path*.

*Proof*. First, we claim that *s* and *t* must be included in 𝒫. The 0^*b*^1 substrings are used as padding to prevent any string in 𝒮 from being matched using portions of two or more vertex labels. Therefore, if *s* or *t* are not included in the inferred path, at least *c* (*n* + 1) + 1 strings from 𝒮 do not occur in *σ*(𝒫), contradicting that *Cost*(𝒫) ≤ *c* (*n* + 1). Hence, the inferred path must contain *s* and *t* and be of the form 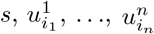 *t* for some *i*_1_, …, *i*_*n*_. Since each edge traversed corresponds to a recombination, the total number of recombinations is *n* + 1. The only way the *Cost*(𝒫) ≤ *c* · (*n* + 1) is if all strings in 𝒮 occur as substrings in *σ*(𝒫). Again, due to the 0^*b*^1 padding in the vertex labels, this can only happen if for all 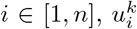 is a vertex in 𝒫for some *k*. Furthermore, because there are *n* vertices in 𝒫that are not *s* or *t*, there must be exactly one such *k* for a given *i*. We conclude that 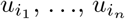 is a Hamiltonian path in *G*_*H*_. □

## 4 Proposed Algorithms

Before developing our integer programming solutions to Problem 1, it is first helpful to define an additional graph representation, which we call as *expanded graph*. In pangenome graphs, multiple haplotype paths share vertices if the sequences are conserved, whereas in the expanded graph, we will split all haplotypes into separate paths (Figures 2A, 2B). The expanded graph enables us to model Problem 1 as a sort of network flow problem. In particular, the inferred path will be reconstructed from a flow of value one in the expanded graph. We will assign weights to edges to account for recombination penalty. Additional constraints will be used to capture how many strings in 𝒮 occur in the resulting inferred path.

**Fig. 2:**
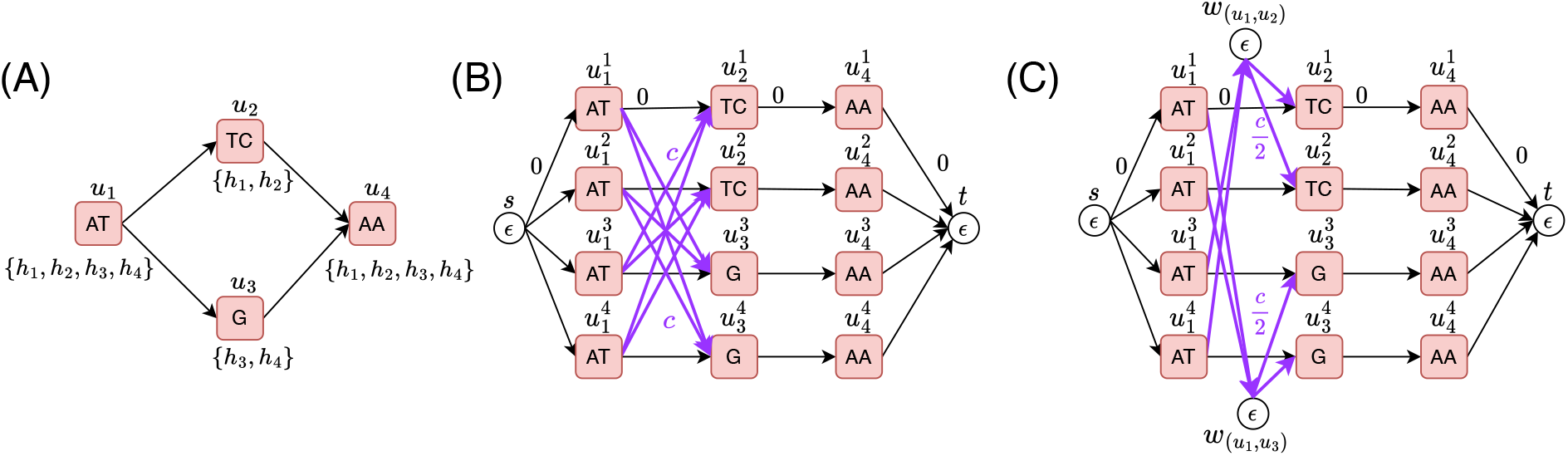
(A) A pangenome graph with four haplotype paths *h*_1_, *h*_2_, *h*_3_ and *h*_4_. Set of haplotype paths passing through a vertex is listed below each vertex. (B) The corresponding expanded graph which includes four disjoint paths, one for each haplotype path. The recombination edges are shown in purple, these edges have a weight of *c*. We consider only the useful recombinations (Lemma 3). The edges which are not recombination edges in the expanded graph have a weight of 0. (C) The corresponding optimized expanded graph.

Lemma 3 allows us to only consider a subset of all possible recombinations in order to find an optimal solution to Problem 1. We call the type of recombination described in Lemma 3 a *useful recombination*.

### Lemma 3.

*There exists an optimal inferred path* 𝒫 = (*a*_1_, …, *a*_*n*_) *for Problem 1 where for all i* ∈ [1, *n*), *a*_*i*_.*h* ≠ *a*_*i*+1_.*h implies vertices a*_*i*_.*u and a*_*i*+1_.*u are not consecutive vertices in haplotype path a*_*i*_.*h*.

*Proof*. Suppose there is an optimal inferred path 𝒫 = (*a*_1_, …, *a*_*n*_) for Problem 1 where for some *a*_*i*_, *a*_*i*_.*h* ≠ *a*_*i*+1_.*h* such that *a*_*i*_.*u* and *a*_*i*+1_.*u* are consecutive vertices in haplotype path *a*_*i*_.*h*. Furthermore, suppose we start with the smallest *i* where this holds. We then change the haplotype path for *a*_*i*+1_ to equal *a*_*i*_.*h*. This does not increase the overall cost, since the number of string 𝒮 occurring in *σ*(𝒫) has not changed, and the number of recombinations either decreases or stays the same. Continuing this process from the next *j > i*, such that *a*_*j*_.*h* ≠ *a*_*j*+1_.*h* and *a*_*j*_.*u* and *a*_*j*+1_.*u* are consecutive vertices in *a*_*j*_.*h*, we achieve an inferred path satisfying the conditions stated in the lemma after at most *n* iterations. □

Next, we present a definition of the expanded graph where we will consider only the useful recombinations. For technical reasons, we preprocess each edge in *E*, splitting it and adding a new vertex labeled with the empty string *ϵ*. Each added vertex inherits the haplotype paths which supported the edge it was formed from. This added step is to prevent recombinations from a haplotype to itself when we build our expanded graph. Now, let *V* = {*u*_1_, …, *u*_*n*_}. For haplotype path 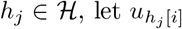, denote the *i*^*th*^ vertex in haplotype path *h*_*j*_. We use *G*_*E*_ = (*V*_*E*_, *E*_*E*_, *σ*_*E*_) to denote the expanded graph. In *G*_*E*_, vertices are string-labeled and edges are weighted. Vertex set *V*_*E*_ is defined as:

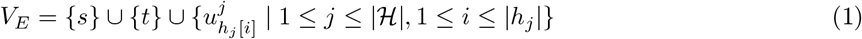

The vertex set contains a source and sink vertex, *s* and *t*, respectively. The vertex set also contains a set of disjoint vertices for each haplotype path in H (Figure 2B). A superscript is used to indicate which haplotype path the vertex is designated to. We refer to the ordered vertex set 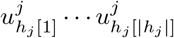 as a haplotype path in *G*_*E*_.

We denote weighted edges in *E*_*E*_ as tuples of the form (*start, end, weight*). The weighted edge set is

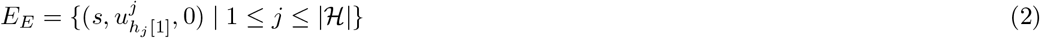

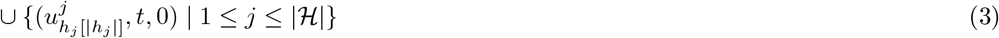

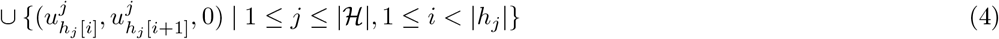

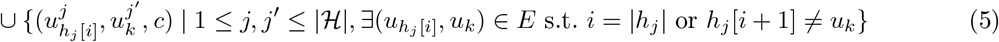

Next, we give some intuition for each line (2)-(5) in the above construction of *E*_*E*_.

(2) Weight 0 edges are created from *s* to the start of each haplotype path in *G*_*E*_.
(3) Weight 0 edges are created from the end of each haplotype path in *G*_*E*_ to *t*.
(4) Weight 0 edges are created between adjacent vertices in each haplotype path. That is, in the path for *h*_*j*_, an edge is created from 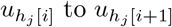
(5) Weight *c* edges are used to represent the useful recombinations described in Lemma 3. We call these *recombination edges*. We use *ϵ* to denote the empty string. The vertex labels are defined as follows:

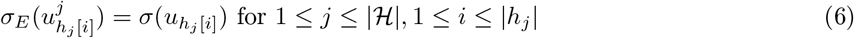

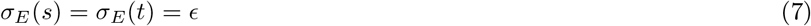
(6) The vertices in a haplotype path are labeled according to the corresponding vertex label in *G*. These labels will be used to identify matches.
(7) The source, sink, do not require vertex labels and are hence labeled with the empty string *ϵ*.

### Optimizing the Expanded Graph

One issue with the above construction is that the number of recombination edges for a given potential recombination can be O(|H|^2^) in the worst case. This occurs because we maintain |*haps*(*ν*)| copies of each vertex *ν* ∈ *V*. For every edge (*u, ν*) ∈ *E* allowing a recombination, we add *O*(|*haps*(*u*)|· |*haps*(*ν*)|) edges to the edge set *E*_*E*_. Since both |*haps*(*u*)| and |*haps*(*ν*)| can be at most |H|, any potential recombination can result in O(|H|^2^) recombination edges in the worst case. We observe this issue in practice as well. An improvement is to represent a recombination by having an intermediate vertex *w*_*e*_ that represents the edge *e ∈ E* allowing for the recombination. We then create an edge to *w*_*e*_ from every vertex in a haplotype path which the recombination would start from, and edges from *w*_*e*_ to every vertex in a haplotype path to which the recombination would lead to (Figure 2C). More formally, the modified vertex set becomes

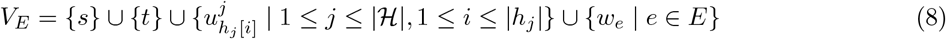

We also replace Line (5) in the construction of *E*_*E*_ with the Lines (9) and (10) as follows: *E*_*E*_ = …

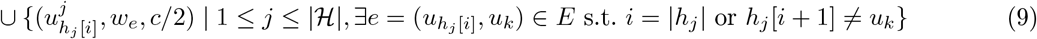

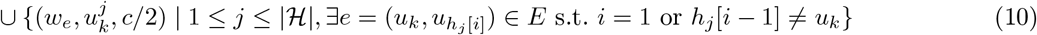

We now call these edges created in Lines (9) and (10) the recombination edges. After creating the edges in *E*_*E*_, we delete any *w*_*e*_ vertex that is isolated in *G*_*E*_. Finally, for any remaining *w*_*e*_ vertices, we define *σ*_*E*_(*w*_*e*_) = *ϵ*. Observe, that the above modification allows for the same set of useful recombinations as our initial expanded graph construction. However, per potential useful recombination, the number of edges remains *O*(|H|) rather than *O*(|H|^2^). Before giving the integer programming solutions, we require one additional definition.

#### Definition 2 (Hits).

*For a string r∈* 𝒮, *assuming* 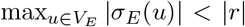, *a path in G*_*E*_, *denoted as an ordered edge set* ((*u, ν*), (*ν, w*), (*w, x*) …, (*y, z*)), *matches r if r* = *σ*_*E*_(*u*)′ · *σ*_*E*_(*ν*) · *σ*_*E*_(*w*) *σ*_*E*_(*x*) ⋯ *σ*_*E*_(*y*) · *σ*_*E*_(*z*)′, *where σ*_*E*_(*u*)′ *is a suffix of σ*_*E*_(*u*) *and σ*_*E*_(*z*)′ *a prefix of σ*_*E*_(*z*). *We use hits*(*r*) *to represent the set of paths matching string r in G*_*E*_.

### 4.1 Integer Linear Programming (ILP) Formulation

We assume that the maximum length of any vertex label is upper bounded by the length of any string in 𝒮, i.e., 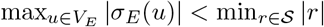. This condition can be easily enforced in the input graph by adjusting the lengths of vertex labels, e.g., by splitting a vertex with a long label into two, while ensuring that the graph’s topology is preserved. We assume min_*r*∈𝒮_ |*r*| *>* 1.

The basis for our solution is to find an *st*-flow with a flow of 1 through the expanded graph *G*_*E*_. Our integer programs will utilize binary decision variable *x*_*uv*_ for each edge. The variable *x*_*uv*_ will take the value 1 if edge (*u, ν*) ∈ *E*_*E*_ is part of the solution flow and 0 otherwise. Because these are binary variables, the flow will always be a path. From the solution path in *G*_*E*_, it is straight forward to recover the corresponding inferred path 𝒫. We use binary decision variable *z*_*r*_ for each string *r* ∈ 𝒮 such that *z*_*r*_ will take the value 1 if the solution flow includes a subpath from *hits*(*r*). We also use variable *z*_*rω*_ for each *ω* ∈ *hits*(*r*), *r* ∈ 𝒮.

Letting *weight*(*u, ν*) denote the weight of an edge (*u, ν*) ∈ *E*_*E*_, our ILP formulation is as follows:

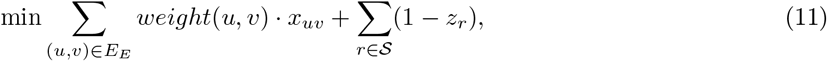

subject to

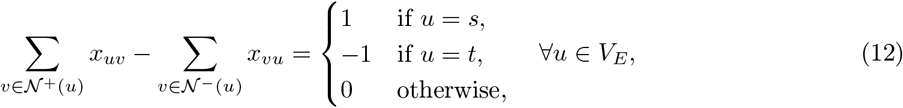

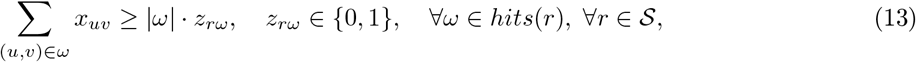

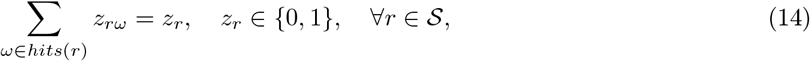

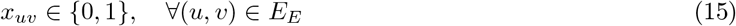

In the ILP formulation, the Objective (11) models *Cost*(𝒫). The summation over *weight*(*u, ν*)·*x*_*uv*_ imposes penalty *c* for each recombination. This is due to the two *c/*2 weighted recombination edges that must traversed when the path switches between haplotype paths in *G*_*E*_ (Figure 2C). In the second summation, the term (1− *z*_*r*_) adds a penalty of 1 to the objective for every *r ∈* 𝒮 where 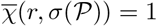 Constraint (12) enforces flow conservation, allowing a unit flow from the source vertex *s* to the sink vertex *t*, ensuring that the ILP formulation selects a single path in the expanded graph.

To explain the function of Constraint (13), termed as linear string-hit constraint and (14), observe that in an optimal solution, whenever possible the variable *z*_*r*_ is set to 1. This is because the term (1− *z*_*r*_) in the objective function adds a penalty of 0 whenever *z*_*r*_ = 1. However, this is only possible when *z*_*rω*_ is equal to 1 for some *ω* ∈ *hits*(*r*). This, in turn, is only possible if Σ _(*u,v*)∈*ω*_ *x*_*uv*_ = |*ω*|, meaning *r* occurs as a substring in the inferred path. Also note that at most one *z*_*rω*_ variable can equal 1 in Constraint (14). Other *z*_*rω*_*′* variables, where *ω, ω*′ ∈ *hits*(*r*) and *ω* ≠ *ω*′, can have a value of 0, even if Σ_(*u,v*)∈*ω*′_ *x*_*uv*_ = |*ω*′|, justifying the use of equality in Constraint (14).

A weakness of the proposed ILP formulation is that the number of string-hit constraints equals the total number of string matches, that is, Σ_r∈𝒮_ *hits*(*r*). We design another formulation with quadratic constraints in which fewer constraints are needed.

### 4.2 Integer Quadratic Programming (IQP) Formulation

In our IQP formulation, Objective (11), and Constraints (12), and (14) and (15) remain unchanged from the ILP formulation. Constraints in (13) are replaced by quadratic constraints defined as

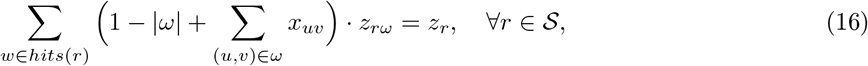

We call Constraint (16) the quadratic string-hit constraint. Again, due to Constraint (14) at most one *z*_*rω*_ variable can be 1. The expression 1− |*ω*| + Σ_(*u,v*) ∈*ω*_ *x*_*uv*_ sums to 1 when the subpath *ω* is contained in the flow. In this case *z*_*r*_ will take the value 1 and no penalty is paid in the objective. Conversely, if some of the edges for *ω* are not in the flow, the expression will sum to ≤ 0. If this is the case for each *ω* ∈ *hits*(*r*), then Constraint (16) can only be satisfied by setting *z*_*r*_ = 0 and *z*_*rω*_ = 0 for each *ω* ∈ *hits*(*r*). Since *z*_*r*_ = 0, a penalty is paid in the objective. The total number of quadratic string-hit constraints is |𝒮|. In our experiments, we observe that IQP formulation solves the problem faster, albeit while requiring more memory.

As a further improvement, we relax the variables *x*_*uv*_ for all (*u, ν*) ∈ *E*_*E*_ to continuous values *x*_*uv*_ *∈* [0, 1] in Constraint (15), following Lemma 4.

#### Lemma 4.

*An optimal solution ϕ*_*cont*_ *to the IQP (or ILP) with relaxed Constraint (15) where variables x*_*uv*_ *lie within the continuous interval* [0, 1] *can be transformed in polynomial time to an optimal solution ϕ satisfying x*_*uv*_ ∈ {0, 1} *for all* (*u, ν*) ∈ *E*_*E*_.

*Proof*. First, observe that *z*_*r*_ = 1 if and only if all edges in some *ω ∈ hits*(*r*) have their corresponding variables set to 1. This follows from Constraints (13) and (16), and the fact that at most one *z*_*rω*_ can be 1 for a given *r*, by Constraint (14).

If *z*_*r*_ = 0 for all *r∈* 𝒮 in *ϕ*_*cont*_, then *ϕ* can be trivially obtained as a single haplotype path in *G*_*E*_ without recombination penalties. In such a case, all edge variables are assigned either 0 or 1.

For the remaining cases, we introduce the following terms:

– *ω ∈ hits*(*r*) is a *used hit-subpath* if *z*_*rω*_ = 1.
– A flow between vertices *u* and *ν* can be decomposed into *uν*-paths each assigned some positive flow and called *flow subpaths*.
– *ω* is the *first used hit-subpath* if there is a flow subpath from vertex *s* to the first vertex of *ω* without passing through another used hit-subpath.
– *ω* is the *last used hit-subpath* if there is a flow subpath from the last vertex of *ω* to vertex *t* without passing through another used hit-subpath.
– *ω* and *ω*′ are *consecutive used hit-subpaths* if there is a flow subpath between them without passing through a third used hit-subpath, where *ω*′ ≠ *ω* and *ω*′ ∈ *hits*(*r*).

Now, if *z*_*r*_ = 1 in *ϕ*_*cont*_ for some *r ∈* 𝒮, there exists a used hit-subpath. We obtain *ϕ* as following. The flow used to reach the first hit-subpath avoids recombination penalties by following a single haplotype path. Similarly, the flow from the end vertex on the last used hit-subpath to *t* avoids recombinations penalties by staying on a single haplotype path. Next, consider two consecutive used hit-subpaths *ω* and *ω*′, with *u* and *ν* as their respective end and start vertices. If *u* and *ν* are on different haplotype paths, any flow subpaths between *u* and *ν* must minimize the recombination penalty. The same minimum recombination cost can be achieved by replacing the potentially multiple fractional flow subpaths with a single path that incurs the same recombination penalty. We can select any flow subpath from *u* to *ν* and assign its edge variables to 1. Edge variables on edges used on the flow from *u* to *ν* and not on this selected path are set to 0. □

## 5 Results

### Implementation Details

We implemented our ILP and IQP solutions in C++ using Gurobi (v11.0.2) solver. We refer to our software as PHI (**P**angenome-based **H**aplotype **I**nference). The user can provide a pangenome reference as either a graph (GFA format) or as a list of phased variants (VCF format). Given short-read or long-read sequencing data of either a haploid or a homozygous genome, PHI outputs the haplotype sequence associated with the optimal inferred path from the graph in FASTA format.

Given a set of reads, we compute (*w, k*) window minimizers [30] for identifying our *hits* (Definition 2). By default, *w* = 25 and *k* = 31. These minimizers correspond to the set 𝒮 in Problem 1. Computing minimizer matches between two strings is faster than computing minimizer matches on a pangenome graph. For this reason, we find minimizer matches between reads and the sequences spelled by all the haplotype paths in the graph. This means *hits*(*r*) includes only those subpaths that are completely contained in some haplotype path in *G*_*E*_ (Definition 2). This restriction to *hits*(*r*) also prevents us from needing to perform the additional edge splitting step described in Section 4.1. We used recombination penalty *c* = 100, this value was chosen empirically. We ran all our experiments on AMD EPYC 7763 processors with 512 GB RAM. We used 32 threads in all experiments.

### Datasets

We evaluated our algorithm by estimating MHC sequences of five haplotypes (APD, DBB, MANN, QBL, SSTO) from homozygous human cell lines. Recently, Houwaart *et al*. [16,32] published complete assemblies of these MHC sequences using long and short-read sequencing. The average length of these assemblies is 4.99 Mbp. We downloaded the five short-read sequencing datasets available from this study. To evaluate our algorithm using varying sequencing coverage, we down-sampled each short-read dataset to obtain coverage of 0.1×, 0.5×, 1×, 2×, 5×, and 10×. We also used the full datasets for evaluation (coverage 12.9 − 18.2×). We used the complete assemblies of five MHC haplotypes as ground-truth to evaluate the accuracy of our estimated sequences. To quantify the accuracy, we measured edit distance between each estimated sequence and the corresponding ground-truth sequence.

We built a haplotype-resolved pangenome graph of 49 complete MHC sequences [19] using Minigraph-Cactus [15]. These sequences were extracted from phased assemblies of 24 diploid human samples [22] and the CHM13 reference [27]. Using Minigraph-Cactus, we obtained the pangenome reference in a VCF format file. We subjected this file to further simplification steps^1^ to ensure compatibility with various tools. We show sequence similarity statistics between the complete MHC assemblies of five haplotypes (APD, DBB, MANN, QBL, SSTO) and the 49 pangenome reference haplotypes in Appendix Table 1.

### Other Methods

We compared PHI with two existing pangenome-based genotyping tools - (i) VG (v1.60) [35] and (ii) PanGenie (v3.1) [9]. VG supports sampling of relevant haplotypes from a pangenome graph by comparing *k*-mer counts in the reads and *k*-mers of a reference haplotype. The selection of haplotypes is done locally in fixed-length non-overlapping blocks. Recombinations may be introduced to create contiguous haplotypes across the blocks. The number of samples can be specified by the user. Accordingly, VG’s haplotype sampling feature can be adapted for haplotype sequence estimation by simply setting the number of desired samples to one. Next, PanGenie supports short-read genotyping using a haplotype-resolved pangenome graph. PanGenie uses a hidden Markov model, which is similar to the standard Li and Stephens model [21]. PanGenie compares *k*-mer counts in the reads with the *k*-mers present in the graph to compute genotype likelihoods. PanGenie exhibited better genotyping accuracy and speed than other genotyping tools [9]. Our sequencing datasets are derived from homozygous cell lines, therefore we ignored the heterozygous genotype calls made by PanGenie (Appendix Table 3). We incorporated PanGenie’s predicted genotypes in the reference sequence to obtain the haplotype sequence. We list our commands to run PHI, VG and PanGenie in Appendix Table 2.

### Genotyping performance

We evaluated PHI, VG and PanGenie methods in their ability to infer the MHC sequences from short read datasets of varying coverage (see Figure 3). Using low coverage datasets (0.1 − 2×), PHI exhibits significantly higher accuracy. VG and PanGenie methods may not be suitable for low-coverage sequencing. For example, the distribution of *k*-mer counts at low coverage can be unreliable. Distinguishing *k*-mers originating from unique versus repetitive regions, as required by PanGenie and VG, is also challenging at low-coverage. Using coverage of 5× or more, the results of VG and PHI are comparable. PanGenie also produces comparable results using full datasets. We note that the integer programming (IQP) approach used in PHI requires more time and memory compared to the methods used in VG and PanGenie. PHI used up to 1.5 hours and 137 GB RAM in a single experiment. In contrast, VG and PanGenie required *<* 5 minutes and *<* 50 GB memory. It may be possible to optimize PHI by incorporating efficient heuristics. We show detailed performance statistics for PHI, including its runtime and memory usage in Appendix Table 4.

**Fig. 3:**
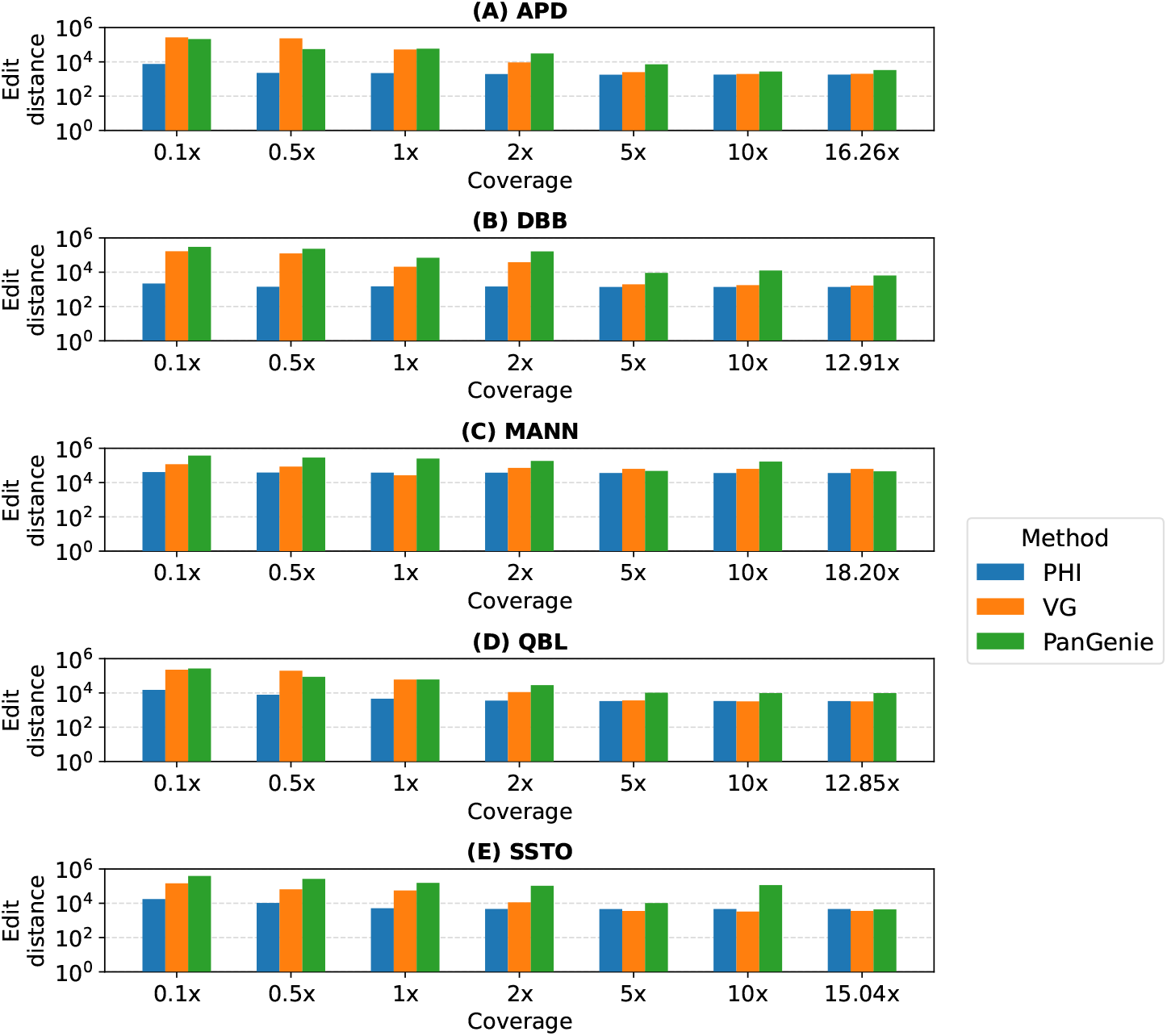
Accuracy of haplotype sequences estimated by PHI, VG and PanGenie using short reads from MHC sequences of five haplotypes (APD, DBB, MANN, QBL, SSTO). The x-axes indicate the coverage of short-read data. The y-axes indicate the edit distance between the estimate haplotype sequence and the ground-truth sequence on a logarithmic scale.

### Effect of our optimizations

In PHI, we implemented both ILP-based and IQP-based solutions to solve the optimization problem. Using either solution, Gurobi solves Problem 1 to optimality. We benchmarked our ILP and IQP solutions to compare their runtime and memory-usage (see Figure 4). On low-coverage datasets (0.1−1×), the runtimes are comparable. At higher coverage, the IQP solution runs faster, which is likely due to fewer string-hit constraints used (Section 4.2). Although, it requires approximately 1.5 times more memory. This may be because Gurobi requires additional storage to handle quadratic constraints. Accordingly, while using PHI, the user can choose between ILP and IQP using a command line argument based on the available memory. If no choice is provided, the IQP solution is used by default. We also evaluated the advantage of relaxing edge variables to continuous values (Lemma 4) by comparing it to another version of our code where we set the edge variables to be discrete. Relaxation of variables deceases runtime of the IQP solution by a factor of 1.6 on average (Appendix Figure 2). Not much effect on the runtime is observed in the ILP solution (Appendix Figure 3).

**Fig. 4:**
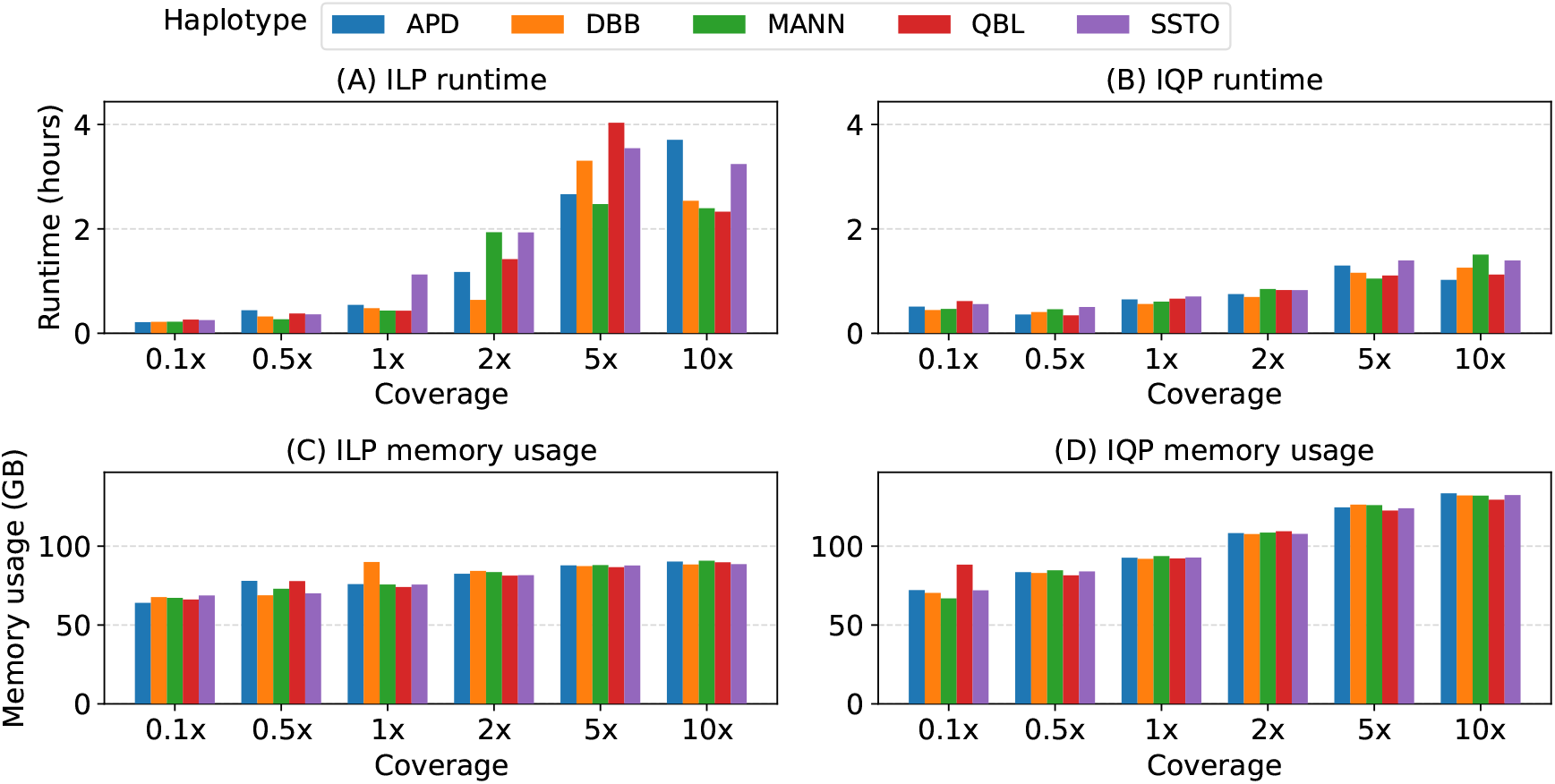
Performance comparison between the ILP and IQP solutions implemented in PHI. We compared their runtime and memory-usage using short-read sequencing datasets sampled from five haplotypes.

### Impact of graph expansion with the addition of more genomes

We evaluated the impact of pangenome graph expansion on PHI’s genotyping accuracy as well as runtime. To do this, we created five versions of our pangenome graph, each containing an increasing number of reference haplotypes, added progressively. The first graph comprises a single diploid sample (chosen randomly from 24 diploid samples) plus CHM13 reference, therefore, it has three reference haplotypes in total. The second graph includes two more diploid samples (chosen randomly from the remaining 23), therefore, it has seven reference haplotypes in total. Similarly, third, fourth and fifth graphs contain 13, 25 and 49 reference haplotypes, respectively. The fifth graph is equivalent to the graph used in previous experiments as well. This results in five different graphs that have 3, 7, 13, 25, and 49 reference haplotypes respectively.

We repeated our experiments with full short-read datasets using these five graphs and present results in Figure 5. We observe that edit distances between the estimated sequences and the ground truth sequences decrease with the increasing number of reference haplotypes. This is expected because more haplotypes are available to choose from when we compute our inferred path in the graph. We also observe an increase in runtime and memory usage. Runtime appears to increase superlinearly and memory appears to increase linearly with the number of reference haplotypes. This is because the size of expanded graph and the number of minimizer matches increase leading to more variables and constraints in our integer program.

**Fig. 5:**
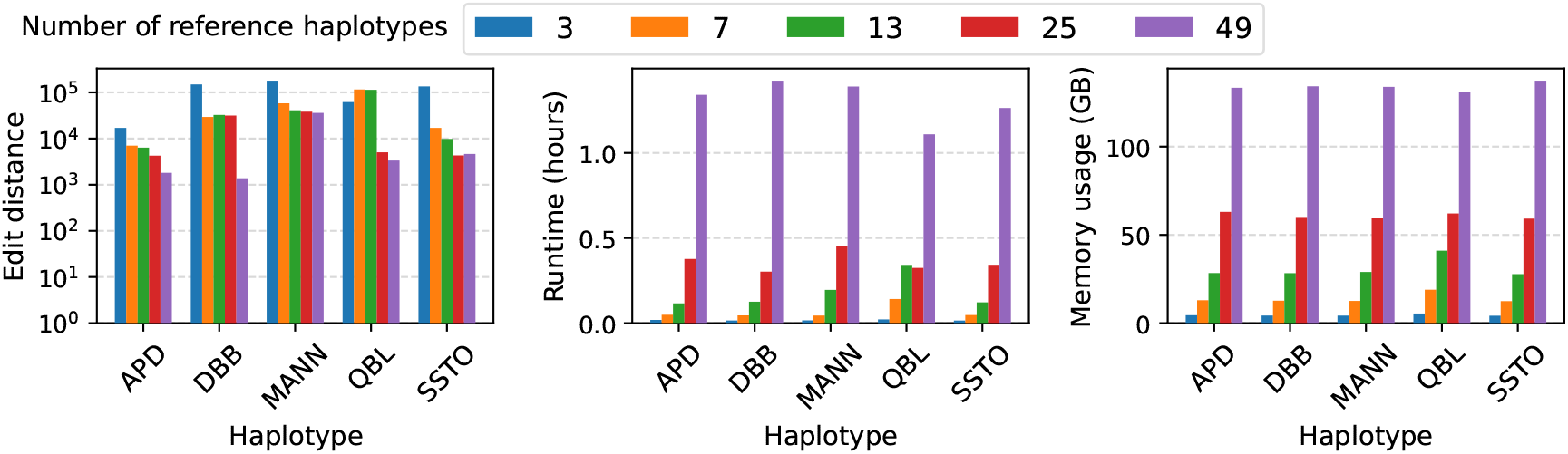
Assessement of PHI’s performance with the increasing number of genomes in pangenome graph. The left figure shows the accuracy in terms of edit distance between the output sequences and ground-truth sequences. The middle and right figure show the runtime and memory-usage respectively.

## 6 Discussion

Genotyping using pangenome graphs is equivalent to finding a walk in the graph that contains the sample’s variants [28]. If the sample is diploid, this becomes equivalent to finding a pair of paths. Drawing inspiration from this idea, we proposed a rigorous framework to infer a path through the graph, such that the sequence spelled by the path is consistent with the sequencing data in terms of the shared *k*-mers between them, while permitting a limited number of recombinations in the path, each incurring a fixed penalty. This optimization problem requires considering all possible paths in the graph. We proved that this problem is NP-Hard and subsequently gave efficient integer programming solutions. As part of our methodology, we introduced the expanded graph data structure on which we could compute an appropriate *st*-flow of 1. Experimental results demonstrate the advantage of the proposed ILP/IQP approaches for accurate genome inference, especially with low-coverage data (coverage 0.1−1×). Thus, our algorithm can facilitate affordable genotyping and association studies of complex and repeat-rich regions of the genome.

Although our approach is currently tailored to haploid samples, it could generalize to diploid samples. This may be accomplished by finding an *st*-flow of 2 through the expanded graph and modifying some constraints. How well this approach genotypes and phases the genome would be interesting to explore. Another limitation of this work is that we do not capture uncertainty. For example, there may be multiple inferred paths with minimum cost. Lastly, pangenome graphs are expected to grow in the number of genomes, therefore, scaling the current approach to a large number of haplotype paths may be important. We leave these extensions to future work.

## Acknowledgements

This research is funded in part by the DBT/Wellcome Trust India Alliance Fellowship (grant number IA/I/23/2/506979), the Intel India Research Fellowship, the National Institutes of Health of the USA (NIHNIAID U01 AI090905), and the Jürgen Manchot Foundation. We utilized computing resources available at the Indian Institute of Science and the U.S. National Energy Research Scientific Computing Center.

## Appendix

**Fig. 1:**
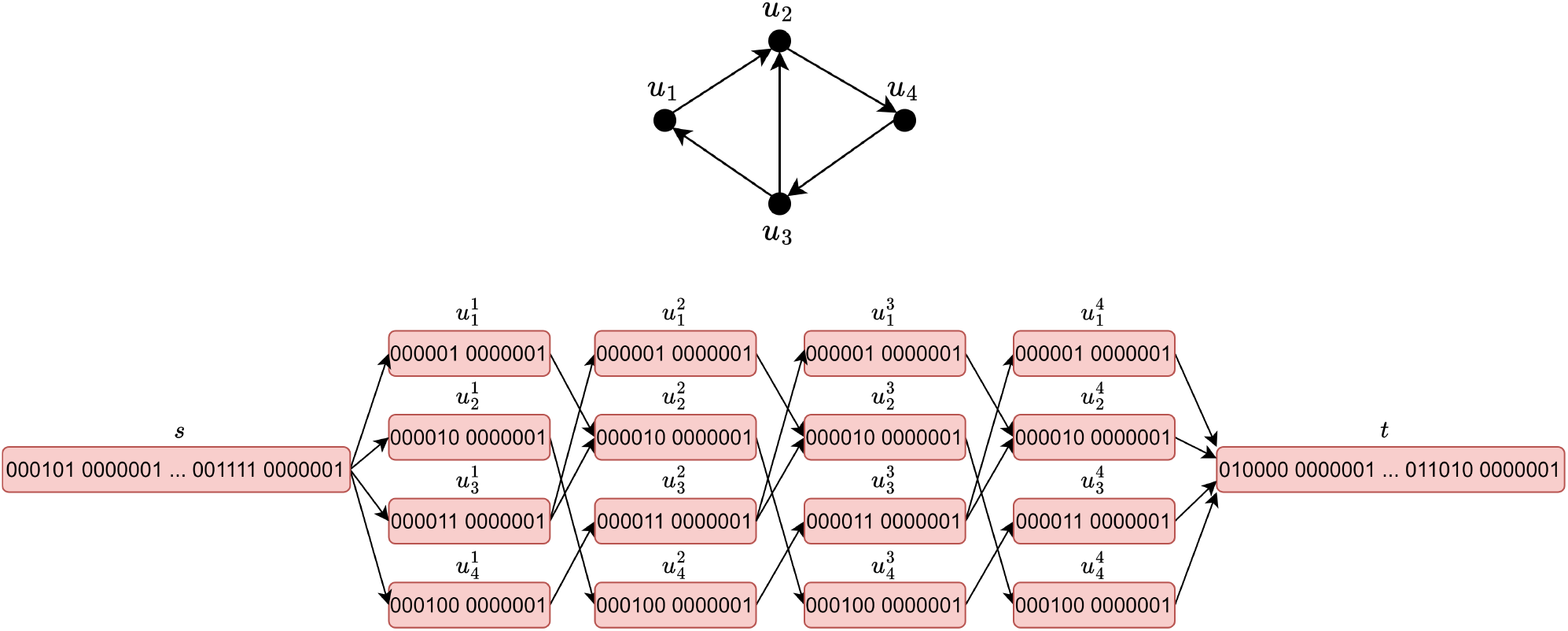
A small example of our reduction from Hamiltonian Path Problem to Problem 1 (Theorem 1). (Top) The starting instance of *G* of Hamiltonian Path Problem. (Bottom) The vertex labeled graph *G*′ constructed from *G*. Here, *n* = 4 and we assume *c* = 2, making *b* = ⌈log_2_(*n* + 2(*c*(*n* + 1) + 1))⌉ + 1 = 6. Each edge is supported by a unique haplotype (not shown). The string set is 𝒮 = {0000010000001, 0000100000001, …, 0110100000001}.

**Table 1:** Additional information about the MHC sequences of five haplotypes (APD, DBB, MANN, QBL, SSTO). We show the length of the complete assembly in the second column. The third and forth columns show edit distance statistics between the assembly and 49 reference haplotypes included in the pangenome reference. In the last two columns, we list the SRA accession numbers and coverage of short-read sequencing datasets.

| Haplotype | Assembly length<br>(Mbp) | Edit distance with<br>pangenome reference haplotypes |  | Short-read data |  |
| --- | --- | --- | --- | --- | --- |
|  |  | Mean | Minimum | SRA Accession | Coverage |
| APD | 4.93 | 146,423 | 37,102 | SRR17272303 | 16.26x |
| DBB | 5.05 | 174,619 | 10,380 | SRR17272302 | 12.91x |
| MANN | 5.03 | 189,464 | 58,168 | SRR17272301 | 18.20x |
| QBL | 4.90 | 159,968 | 72,293 | SRR17272300 | 12.85x |
| SSTO | 5.05 | 161,044 | 35,583 | SRR17272299 | 15.04x |

**Table 2:**
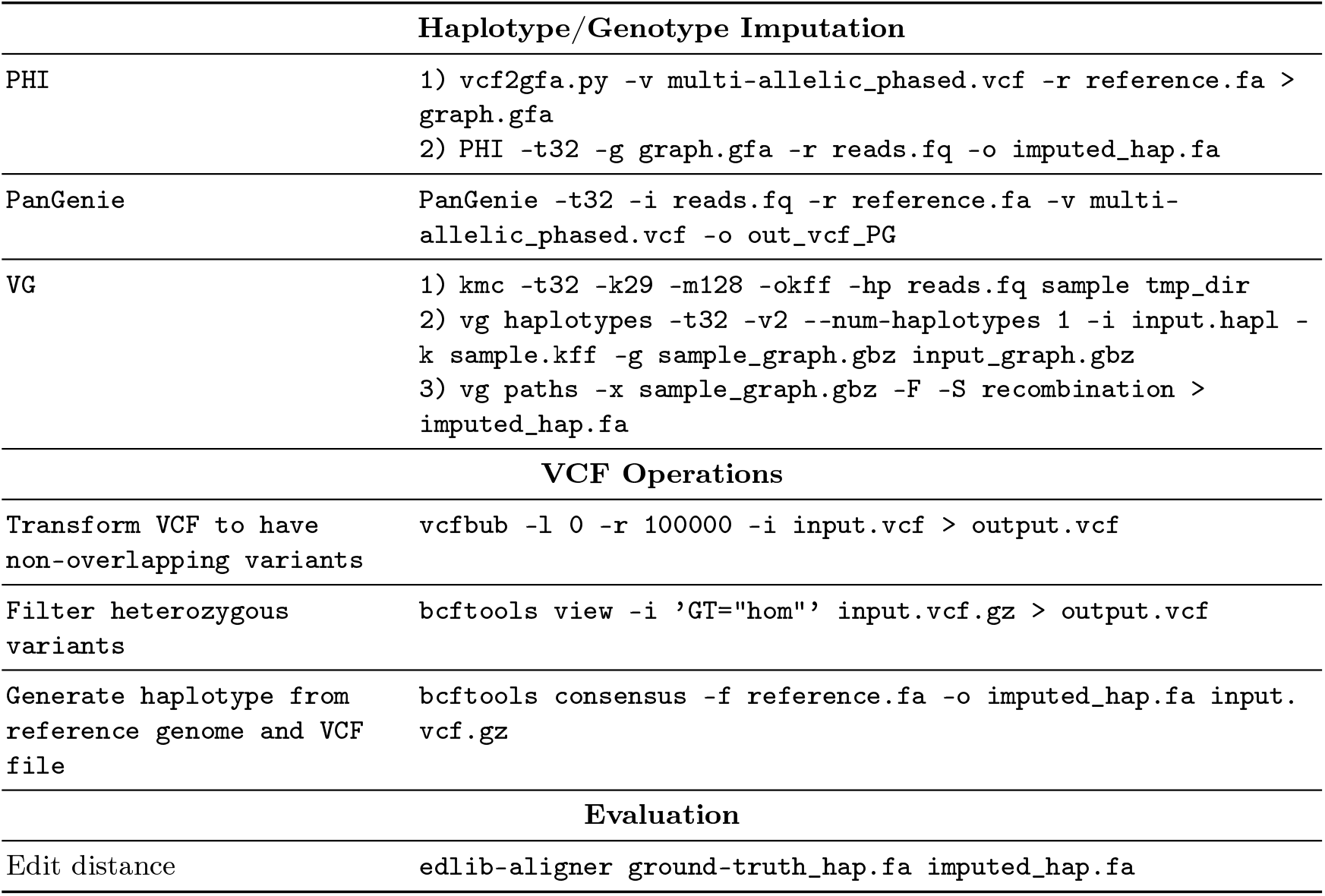
Commands used for running various tools.

| **Haplotype/Genotype Imputation** |  |
| --- | --- |
| PHI | 1) vcf2gfa.py -v multi-allelic_phased.vcf -r reference.fa > graph.gfa<br>2) PHI -t32 -g graph.gfa -r reads.fq -o imputed_hap.fa |
| PanGenie | PanGenie -t32 -i reads.fq -r reference.fa -v multi-allelic_phased.vcf -o out_vcf_PG |
| VG | 1) kmc -t32 -k29 -m128 -okff -hp reads.fq sample tmp_dir<br>2) vg haplotypes -t32 -v2 --num-haplotypes 1 -i input.hapl -k sample.kff -g sample_graph.gbz input_graph.gbz<br>3) vg paths -x sample_graph.gbz -F -S recombination > imputed_hap.fa |
| **VCF Operations** |  |
| Transform VCF to have non-overlapping variants | vcfbub -l 0 -r 100000 -i input.vcf > output.vcf |
| Filter heterozygous variants | bcftools view -i 'GT="hom"' input.vcf.gz > output.vcf |
| Generate haplotype from reference genome and VCF file | bcftools consensus -f reference.fa -o imputed_hap.fa input.vcf.gz |
| **Evaluation** |  |
| Edit distance | edlib-aligner ground-truth_hap.fa imputed_hap.fa |

**Table 3:** Count of homozygous and heterzygous genotype calls made by PanGenie. In our benchmark, we excluded the heterozygous calls because the sequencing datasets were derived from homozygous cell lines.

| Coverage | APD |  | DBB |  | MANN |  | QBL |  | SSTO |  |
| --- | --- | --- | --- | --- | --- | --- | --- | --- | --- | --- |
|  | Hom | Het | Hom | Het | Hom | Het | Hom | Het | Hom | Het |
| 0.1x | 52,816 | 6,245 | 51,435 | 7,626 | 52,452 | 6,609 | 53,707 | 5,354 | 53,893 | 5,168 |
| 0.5x | 56,249 | 2,812 | 55,845 | 3,216 | 56,258 | 2,803 | 56,447 | 2,614 | 56,064 | 2,997 |
| 1x | 57,448 | 1,613 | 57,010 | 2,051 | 57,064 | 1,997 | 57,224 | 1,837 | 57,099 | 1,962 |
| 2x | 58,201 | 860 | 57,948 | 1,113 | 58,334 | 727 | 58,101 | 960 | 58,397 | 664 |
| 5x | 58,552 | 509 | 58,382 | 679 | 58,601 | 460 | 58,340 | 721 | 58,228 | 833 |
| 10x | 58,533 | 528 | 58,478 | 583 | 58,188 | 873 | 58,343 | 718 | 58,337 | 724 |
| Complete data | 58,647 | 414 | 58,457 | 604 | 58,592 | 469 | 58,457 | 604 | 58,521 | 540 |

**Table 4:** We report additional performance statistics for PHI on all our datasets. We specify the number of recombinations used in the solution in the second column. Next, we mention the runtime and memory usage of PHI. In the fifth and the sixth columns, we specify edit distance and alignment identity between the output MHC sequence and the ground-truth sequence. Alignment identify is defined as the ratio of the number of character matches divided by the length of the alignment. In the last three columns, we give statistics about the minimizers computed from sequencing reads. We give the count of distinct minimizers observed in the read set. A fraction of minimizers would be absent from the graph, and some fraction would be present in all reference haplotypes, making them ‘uninformative’. The matches of only the remaining fraction minimizers are useful while solving the optimization problem.

| Coverage | Recombinations | Time<br>(s) | Memory<br>(GB) | Edit<br>distance | Alignment<br>identity (%) | Minimizers<br>(Reads) | Minimizers<br>% Absent \| % Uninformative |  |
| --- | --- | --- | --- | --- | --- | --- | --- | --- |
| Haplotype: APD |  |  |  |  |  |  |  |  |
| 0.1× | 3 | 1840 | 72 | 7551 | 99.85 | 33248 | 36.33 | 43.12 |
| 0.5× | 7 | 1294 | 84 | 2272 | 99.95 | 156209 | 37.90 | 41.42 |
| 1× | 7 | 2338 | 93 | 2220 | 99.95 | 289795 | 41.46 | 39.19 |
| 2× | 9 | 2702 | 108 | 1948 | 99.96 | 508720 | 46.47 | 35.84 |
| 5× | 10 | 4671 | 125 | 1779 | 99.96 | 984355 | 59.39 | 27.05 |
| 10× | 10 | 3683 | 134 | 1810 | 99.96 | 1599325 | 72.22 | 18.33 |
| 16.26× | 10 | 4536 | 134 | 1810 | 99.96 | 2288126 | 80.17 | 13.00 |
| Haplotype: DBB |  |  |  |  |  |  |  |  |
| 0.1× | 2 | 1604 | 70 | 2191 | 99.96 | 33901 | 37.28 | 41.78 |
| 0.5× | 4 | 1467 | 83 | 1415 | 99.97 | 157510 | 39.66 | 39.60 |
| 1× | 4 | 2022 | 92 | 1496 | 99.97 | 293996 | 42.54 | 37.84 |
| 2× | 4 | 2502 | 108 | 1472 | 99.97 | 518085 | 47.59 | 34.28 |
| 5× | 4 | 4175 | 126 | 1385 | 99.97 | 1015730 | 60.37 | 25.75 |
| 10× | 4 | 4525 | 132 | 1377 | 99.97 | 1660305 | 72.79 | 17.55 |
| 12.91× | 4 | 4743 | 135 | 1377 | 99.97 | 2028107 | 77.31 | 14.58 |
| Haplotype: MANN |  |  |  |  |  |  |  |  |
| 0.1× | 3 | 1680 | 67 | 41028 | 99.19 | 33614 | 34.31 | 43.07 |
| 0.5× | 7 | 1658 | 85 | 38379 | 99.24 | 153933 | 36.66 | 41.50 |
| 1× | 8 | 2183 | 94 | 37898 | 99.25 | 288713 | 39.33 | 39.76 |
| 2× | 9 | 3054 | 109 | 37728 | 99.25 | 502336 | 44.89 | 36.22 |
| 5× | 12 | 3774 | 126 | 36263 | 99.28 | 964364 | 57.71 | 27.55 |
| 10× | 14 | 5426 | 132 | 35941 | 99.29 | 1553694 | 70.85 | 18.86 |
| 18.20× | 14 | 4843 | 134 | 35940 | 99.29 | 2450244 | 81.06 | 12.15 |
| Haplotype: QBL |  |  |  |  |  |  |  |  |
| 0.1× | 3 | 2222 | 88 | 15062 | 99.69 | 32464 | 35.13 | 43.05 |
| 0.5× | 9 | 1236 | 81 | 7829 | 99.84 | 153818 | 37.47 | 41.77 |
| 1× | 10 | 2388 | 92 | 4610 | 99.91 | 284587 | 39.92 | 40.35 |
| 2× | 14 | 2981 | 109 | 3561 | 99.93 | 502087 | 46.98 | 36.14 |
| 5× | 17 | 3986 | 123 | 3349 | 99.93 | 966151 | 58.80 | 27.40 |
| 10× | 17 | 4049 | 129 | 3356 | 99.93 | 1566636 | 71.76 | 18.63 |
| 12.85× | 17 | 4113 | 131 | 3343 | 99.93 | 1862566 | 75.90 | 15.84 |
| Haplotype: SSTO |  |  |  |  |  |  |  |  |
| 0.1× | 2 | 2013 | 72 | 17626 | 99.65 | 33792 | 36.06 | 41.98 |
| 0.5× | 12 | 1812 | 84 | 10471 | 99.79 | 156473 | 37.60 | 41.12 |
| 1× | 20 | 2536 | 93 | 5150 | 99.90 | 291484 | 41.05 | 38.59 |
| 2× | 24 | 2977 | 108 | 4671 | 99.91 | 513683 | 46.50 | 35.02 |
| 5× | 24 | 5023 | 124 | 4611 | 99.91 | 992511 | 59.01 | 26.68 |
| 10× | 24 | 5021 | 132 | 4634 | 99.91 | 1609715 | 71.88 | 18.16 |
| 15.04× | 24 | 4499 | 137 | 4637 | 99.91 | 2206289 | 79.07 | 13.44 |

**Fig. 2:**
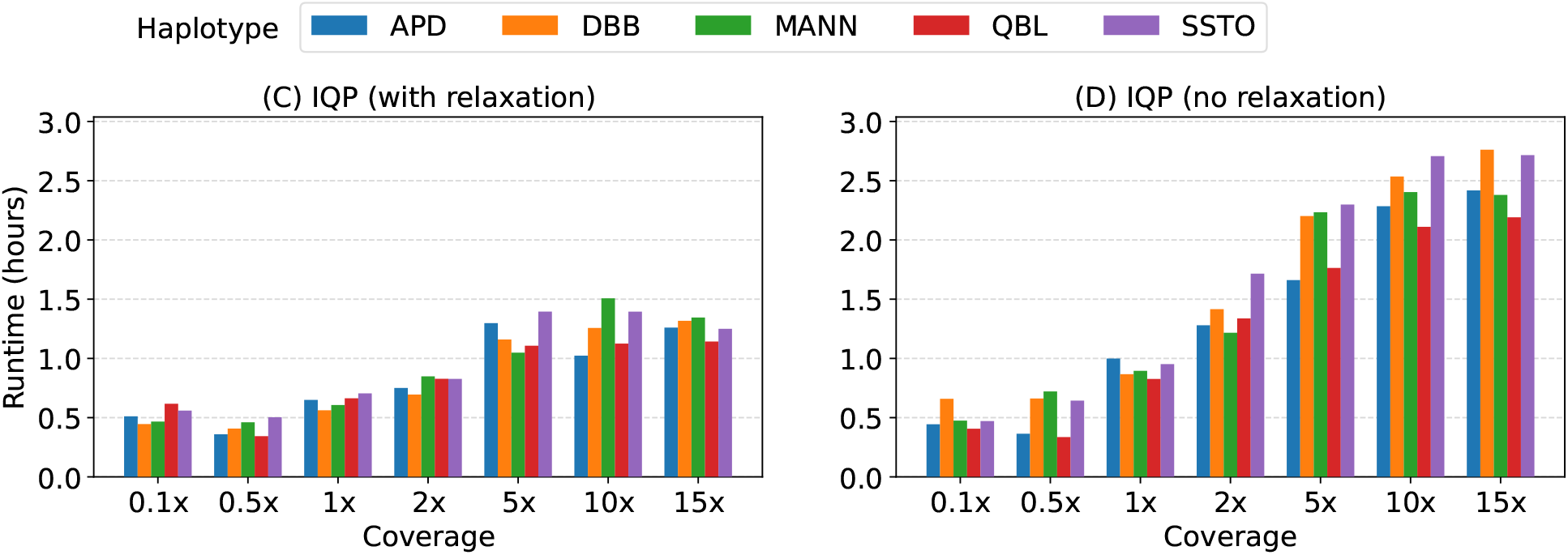
Evaluation of the performance of the IQP method with and without relaxation of the binary edge variables *x*_*uv*_. We compared runtime using various short-read datasets.

**Fig. 3:**
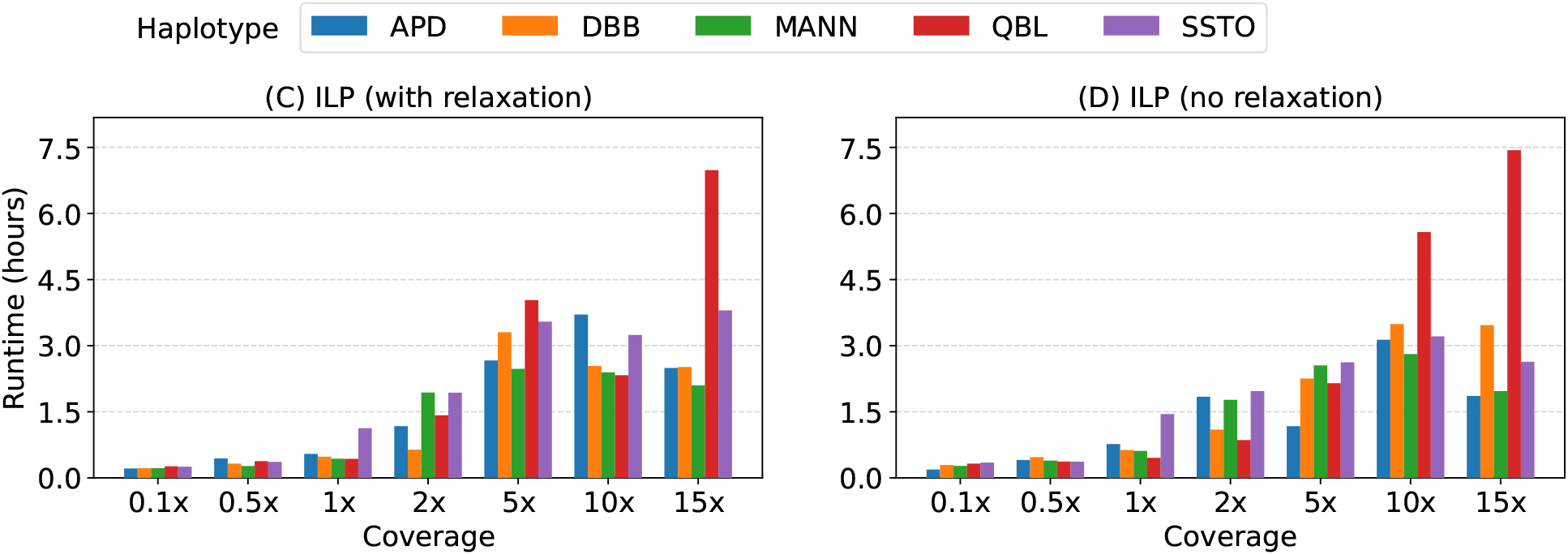
Evaluation of the performance of the ILP method with and without relaxation of the binary edge variables *x*_*uv*_. We compared runtime using various short-read datasets.

## Footnotes

1 https://github.com/eblerjana/genotyping-pipelines/tree/main/prepare-vcf-MC

